# Nanopore Sequencing Reveals High-Resolution Structural Variation in the Cancer Genome

**DOI:** 10.1101/209718

**Authors:** Liang Gong, Chee-Hong Wong, Wei-Chung Cheng, Harianto Tjong, Francesca Menghi, Chew Yee Ngan, Edison T. Liu, Chia-Lin Wei

**Author notes:** These authors contributed equally to this work (Liang Gong, Chee-Hong Wong). Correspondence should be addressed to C.-L.W.

## Abstract

Acquired genomic structural variants (SVs) are major hallmarks of the cancer genome. Their complexity has been challenging to reconstruct from short-read sequencing data. Here, we exploit the long-read sequencing capability of the nanopore platform using our customized pipeline, *Picky*, to reveal SVs of diverse architecture in a breast cancer model. From modest sequencing coverage, we identified the full spectrum of SVs with superior specificity and sensitivity relative to short-read analyses and uncovered repetitive DNA as the major source of variation. Examination of the genome-wide breakpoints at nucleotide-resolution uncovered micro-insertions as the common structural features associated with SVs. Breakpoint density across the genome is associated with propensity for inter-chromosomal connectivity and transcriptional regulation. Furthermore, an over-representation of reciprocal translocations from chromosomal double-crossovers was observed through phased SVs. The comprehensive characterization of SVs using the robust long-read sequencing approach in cancer cohorts will facilitate strategies to monitor genome stability during tumor evolution and improve therapeutic intervention.

## Introduction

Genomic structural variation is prevalent in the human genome^1^ and includes deletions, insertions, duplications, inversions, and translocations. Collectively, these structural variants (SVs) account for a significant portion of genome heterogeneity between individuals^2^ and human populations^3^. The complexity and diversity of genome structure changes have been shown to influence genome evolution and genetic diversity among populations, while specific SVs are associated with diseases susceptibility^4-6^. Many cancer genomes have been found to harbor significant structural variation, and specific SVs are considered to be instrumental in promoting tumor progression by disrupting gene structures, dysregulating gene expression, creating fusing transcription units or increasing gene copy number^7-9^. The detection of specific SVs can be used as the basis for tumor classification and potentially of prognostic value for tumor severity and therapeutic response^7-10^. However, despite the prevalence of SVs and their particular relevance to cancer, the molecular organization of various SV classes and the mechanisms that generate them are not well understood. This is in large part due to the inability of current technologies to uncover the full spectrum of SVs with high specificity at nucleotide-level resolution.

Advances in sequencing technology coupled with improvements in computational algorithms have greatly enhanced our understanding of the abundance, diversity, and molecular features of SVs across human populations^3^ and disease^4, 6^. However, current sequencing approaches that generate high coverage, paired-end short-read sequencing data, combined with split read mapping methods lack the precision and sensitivity necessary for the detection of many types of SVs, particularly in regions of repetitive sequence^11^. Specifically, paired-end short reads are not sufficiently sensitive to detect small SVs, require a high depth of sequencing coverage to achieve high specificity, and lack the nucleotide-level of detail for analysis of the breakpoints that flank SVs. They are also unable to decipher complex SV patterns or provide haplotype phase of SVs in diploid genomes. Thus, there is an ongoing effort to develop effective long-read sequencing approaches for the characterization of SVs^12, 13^.

Recent progress in nanopore single-molecule sequencing offers to extend sequencing read length and throughput, features that would facilitate detection and analysis of SVs in large complex genomes^14^. The nanopore platform sequences individual DNA molecules by passing them through nanometer-scaled pores and measuring changes in the electric current across the pore caused by the transiting nucleotides. These current changes are computationally deconvoluted to reveal the identity of the bases^15, 16^. Unlike sequencing-by-synthesis technology platforms, nanopore sequencing can generate reads of 10–100 Kb in real-time^17, 18^. Thus, nanopore sequencing offers the speed, read length and throughput necessary for comprehensive and unbiased SV profiling and would be particularly useful for resolving complex structural rearrangements in cancer genomes.

Here we demonstrate the utility of nanopore long-read sequencing to detect the full spectrum of SVs and characterize their genomic breakpoints with high specificity and sensitivity. To exploit the value of long reads in SV detection, we developed a computational analysis pipeline, named *Picky*, which optimally performs read alignments and logically defines the full spectrum of SVs, including complex SVs enriched in repetitive DNA elements. From less than 8 Gb of sequencing data, we uncovered 34,100 SVs and 66,660 breakpoints in a well-studied breast cancer cell line. We found that the phasing of SVs suggests frequent reciprocal inter-chromosomal translocations. From analysis of SV breakpoints at nucleotide-resolution, we revealed that micro-insertions are prevalent within break junctions and that breakpoint density is associated with the frequency of interactions between chromatin territories and transcriptional activity. Our results not only establish that long reads from nanopore sequencing enable robust and highly accurate SV detection but also reveal new insights into architecture of SVs that suggest mechanisms for their generation.

## Results

### Read length, throughput and accuracy of nanopore sequencing

To assess the utility of the nanopore platform for detecting SVs, we used it to sequence the genome of the breast cancer cell line HCC1187^19^, a model for triple-negative breast cancer (TNBC) whose genome harbors extensive structural variation that has been previously characterized at the molecular level by paired-end short-read whole genome sequencing^10, 20^. Thus, we can compare the results of our long-read sequencing analysis to this previous data set. Nanopore sequencing libraries were prepared from fragmented high-molecular-weight genomic DNA and subjected for sequencing on MinION instruments (**Methods**). Sequencing of the single strand templates of the DNA fragments was performed to generate 1D data sets, while sequencing of both the template and complement strands, enabled through their covalent bridging by a hairpin adaptor, was performed to generate 2D data sets (**Fig. 1a**). The 2D data sets improve sequence accuracy by aligning the template and complement sequences and resolving any ambiguous base calls between them in the final read output.

**Figure 1.**
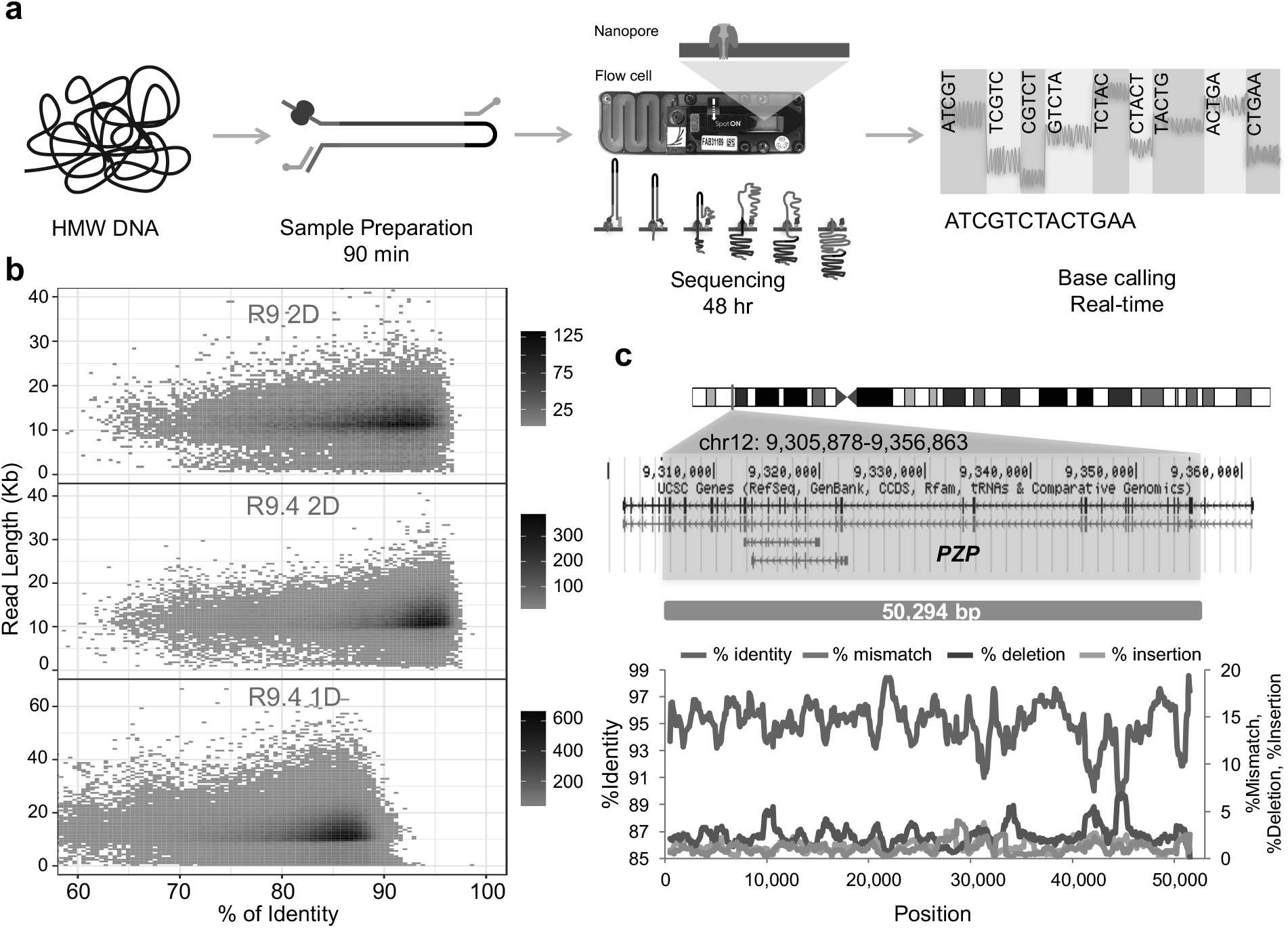
Nanopore single-molecule sequencing. (**a**) Schematic overview of nanopore sequencing process. The ends of double-stranded genomic DNA (blue and red) are ligated with both leader adaptor (Y adaptor, grey) bound a motor protein (purple), hairpin adaptor (black) and annealed to the tethering oligos (green and orange). Sequencing starts from the 5’ end of the leader adaptor. The motor protein unwinds the dsDNA allowing single-stranded DNA to pass through the nanopore (dark cyan). The hairpin adaptor covalently links the template and complement DNA strands, which allows the complementary strand of DNA to pass through the nanopore to 2D reads. Sensors record the current perturbation as ssDNA passes through the nanopore. Each event is base-called to the corresponding 5-mer by a statistical model. (**b**) Accuracy of nanopore reads from different versions of chemistries and run protocols. The percentages of identity among different read length distribution were shown for 2D reads in R9 (250 bases/sec), R9.4 (250 bases/sec) and 1D read (R9.4, 450 bases/sec). **(c**) A 50 Kb nanopore 2D read aligns to reference human genome (chr12: 9,305,878-9,356,863) with 95% average identity. The major error profiles are shown at the below.

Using the R9 and R9.4 MinION-compatible chemistries, we achieved sequencing speeds of 250–450 bases/sec. Although substantial variations in flow cell quality were observed, significant improvement in total yield was associated with increasing sequencing speed (**Supplementary Fig. 1a** and **Supplementary Table 1**). These sequences were aligned onto the reference genome assembly using the genome alignment tool LAST^21^ and the alignment quality was used to assess the degree of accuracy and identify base-calling errors for the different protocols and chemistries. As expected, the accuracy of the 2D reads was on average higher (94%) than that of the 1D reads (86%). Over 70% of the reads from the R9.4 chemistry aligned to reference genome with ≥ 90% identity compared to only 21% from the R9 chemistry (**Fig. 1b**). Based on these data, we used 2D reads for our subsequent analyses (**Supplementary Fig. 1b**).

From 7.9 Gb of the aligned sequence data, we obtained 2.5X average genome coverage (based on a haploid genome size) (**Supplementary Fig. 1c**). Compared to paired-end short reads at equivalent sequencing depth, nanopore long reads extended further into repetitive sequence regions of the genome and covered more of the genome (82% vs. 77%) (**Supplementary Fig. 2a**). For example, a 14.7-Kb nanopore read extended into a region that is rich in short interspersed nuclear elements/long interspersed nuclear elements (SINEs/LINEs) (chr1: 25,732,083-25,747,923), a gap in the coverage of an ultra-high depth (185 Gb, 60X) of short-read data (**Supplementary Fig. 2b**). To determine if nanopore sequencing has any obvious length bias, we subjected DNA templates of different sizes (3–4 Kb and 12 Kb) to sequencing. We found that the resulting read length distributions matched well with the input DNA fragment sizes (**Supplementary Fig. 3a,b**) and the total amounts of bases sequenced were equivalent (**Supplementary Table 1**). When compared with the read length distribution from the established PacBio SMRT long-read sequencing platform, using identical preparation of 12 Kb DNA templates, nanopore exhibited less bias in read length (**Supplementary Fig. 3c**). One of the longest 2D reads we obtained was 50 Kb (i.e. totaling over 100 Kb for sequencing template and complement strands). This read aligned to the *PZP* gene region (chr12: 9,305,878-9,356,863) at an average of 95% identity (**Fig. 1c**), suggesting that nanopore is capable of generating read length beyond 100 Kb. Taken together, the ability to generate long reads at gigabase output with high accuracy indicates that nanopore sequencing can be adopted to effectively analyze structural variation in cancer genomes.

### Picky: an analysis pipeline developed specifically to detect SVs using long read data

To support detection of SVs using long read data, we devised an end-to-end analysis pipeline named “*Picky”*. *Picky* analyzes long reads in three consecutive steps: read alignment to a reference genome, optimal alignment merge/selection, and SV classification (**Fig. 2a**). First, *Picky* adopts LAST^21^ to perform genome alignment. To accommodate errors in nanopore reads and achieve higher sensitivity, we adopted the scoring scheme used in the megaBLAST alignment parameters (href="https://www.ncbi.nlm.nih.gov/books/NBK279678/). In the second step, alignments for each read were evaluated for their quality and spurious alignments were filtered out based on poor alignment score or low percentage identity. Next, alignments for different segments of a long read were picked and merged. We applied a greedy seed-and-extension algorithm to stitch segments together and then combined segments that maximized coverage for each long read. Only reads with >70% genome alignments across their total length were used for further analysis. Reads having multiple segments that each aligned to unique but non-contiguous genomic locations of the reference genome were defined as ‘split reads’. Based on the order and the distance between the non-contiguous alignments, *Picky* assigned split reads into seven classes of SVs: inversion (INV): segments aligned to the same chromosome but in different orientations; translocation (TLC): segments aligned to different chromosomes; tandem duplication (TD): segments contain a complete duplicated region (TDC) or only span across a duplication junction (TDJ); simple insertion (INS) or deletion (DEL), segments correspond to the same chromosomal region in the same orientation but either flank a sequence that does not match the reference genome or lack an intervening sequence observed in the reference genome, respectively; and (INDEL), segments indicate both INS and DEL for the same split read (**Methods**).

**Figure 2.**
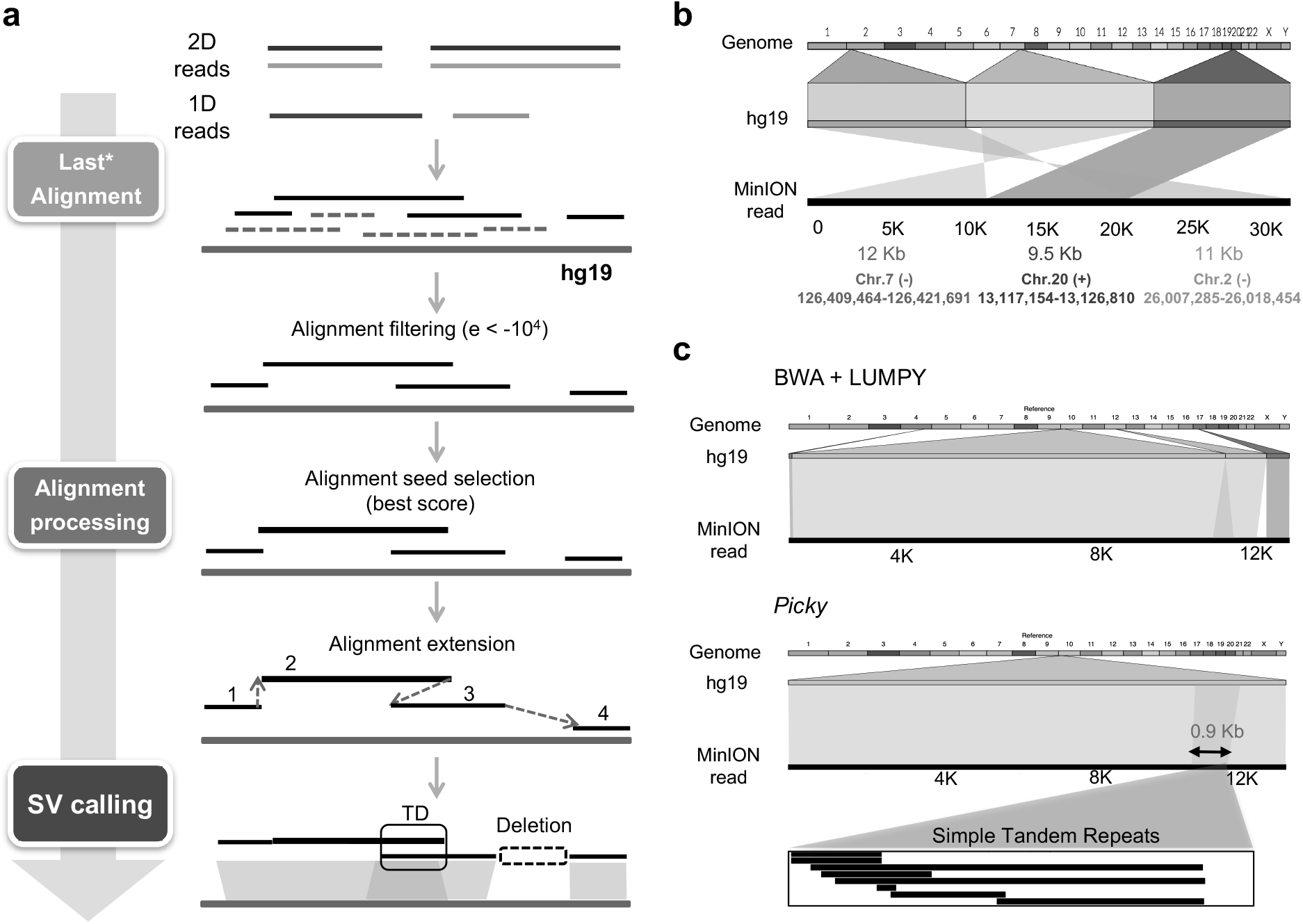
A customized SV pipeline designed for long-read SV analysis. (**a**) Overview of the *Picky* pipeline. The major modules include read alignment, filtering, optimal alignment selection, extension and SV classification (**Methods**). Blue and light blue lines indicate the template and the complementary reads from nanopore 2D reads, while brown and light brown lines indicate different strands of nanopore 1D reads. * indicates minor modification. **(b**) A complex SV of two translocations detected by *Picky* from a 32.5 Kb nanopore read. **(c**) *Picky* accurately defines a short TD. A 12.9 Kb nanopore read aligns to reference genome as two overlapping segments of 11.4 Kb and 2.4 Kb with 86% and 83% identities (e-value = 0). This TD was misclassified as a translocation by the short read aligner BWA and SV caller LUMPY. The alignments were visualized by Ribbon (https://github.com/MariaNattestad/ribbon).

We applied *Picky* to 796,029 2D reads to detect and classify SVs in the HCC1187 genome (**Fig. 3a**). From 53,701 split reads, *Picky* detected a total of 34,100 unique SVs and their corresponding 66,660 breakpoints, respectively, and classified them into 220 inversions, 1,911 translocations, 3,567 tandem duplications and 28,402 insertions, deletions and INDEL (**Supplementary Tables 2** and **3**). Consistent with the notion that longer reads are more likely to capture breakpoints and SVs, the percentage of reads containing breakpoints (i.e. split reads) positively correlated with DNA read lengths, from 2–6% in 2.5–5.5 Kb reads to 13–14% in 12 Kb reads (**Supplementary Fig. 4**). Furthermore, 4% (2,177 of 53,701) of split reads contained more than two breakpoints, which indicates the presence of multiple SVs on the same chromosomes. This phasing information is uniquely provided by long read sequencing. As an example, a nanopore read of 32.5 Kb consisted of three consecutive segments of 12, 9.5 and 11 Kb that each aligned to a different chromosome (chromosome 7, 20 and 2) with high identity (> 97%) (**Fig. 2b**). This read captured two translocations that fused the 5’ exons of *GRM8* to the introns from *SPTLC3* and *ASXL2,* resulted in a non-functional transcript. Through examination of adjacent SVs for which we could glean phasing information via multiple breakpoints within long reads, we found 67 co-occurring translocations (TLC-TLC), representing an enrichment of 3.09 (obs/exp) over the background (**Supplementary Fig. 5a,b** and **Methods**). Interestingly, 37% of dual-translocation events (25 of 67) were reciprocal (i.e. chromosome A-B-A), suggesting that double-crossovers between two non-homologous chromosomes could be a common mode to generate translocation. Therefore, *Picky*-mediated analysis of nanopore long reads can resolve complex chromosomal rearrangements and delineate haplotype specific SVs.

**Figure 3.**
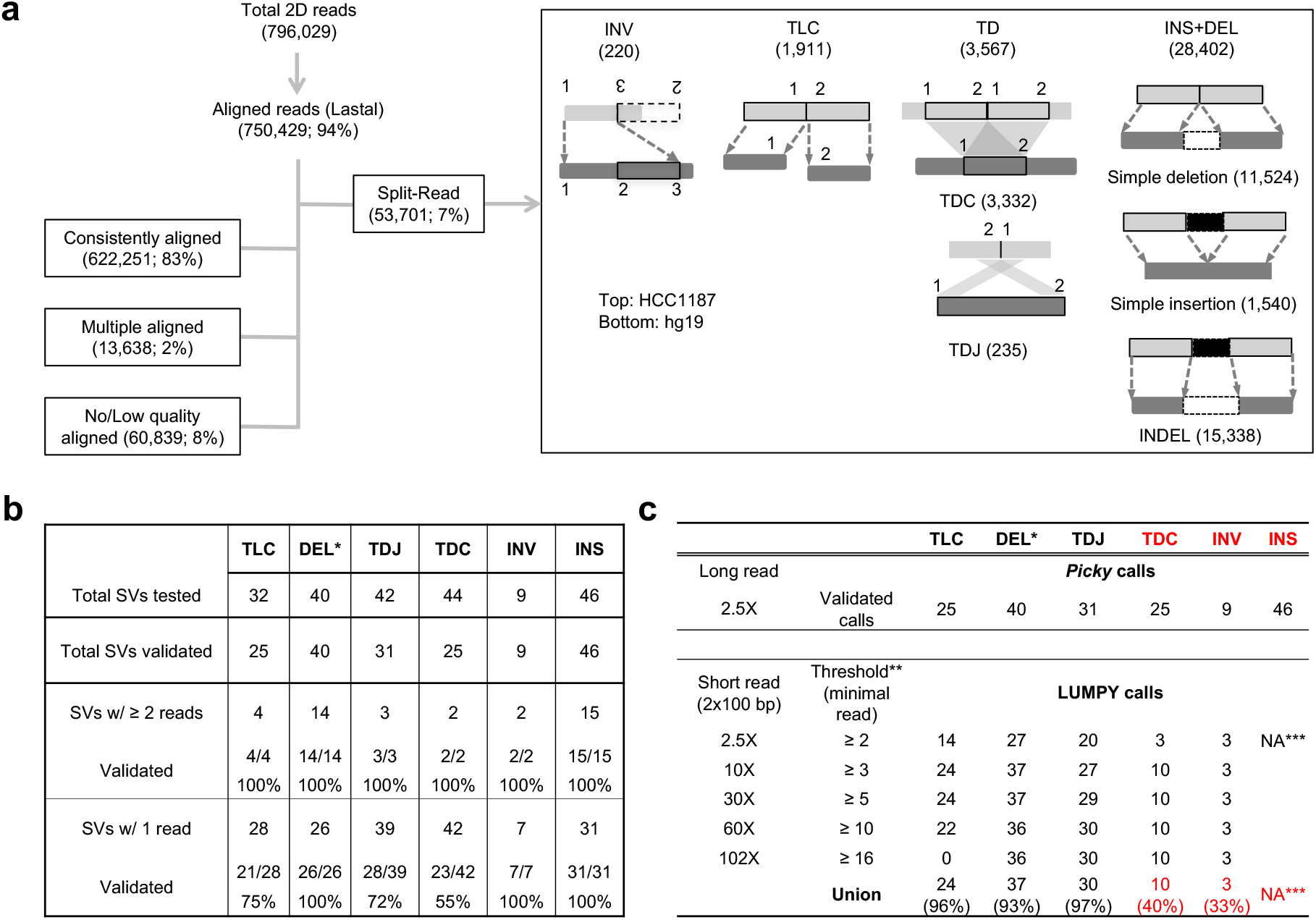
Summary of read alignment, SV detection and validation. (**a**) Read alignment summary and different classes of SVs detected by algorithmic design of *Picky*. **(b**) Numbers of structural variants confirmed by PCR from different SV classes. (**c**) Numbers of SVs called by LUMPY from different depth of short-read data. *: simple deletion and DEL in INDEL. **: thresholds used in SV calling by LUMPY (**Methods**). ***: not called by standard LUMPY pipeline.

In contrast to short read SV callers such as LUMPY, which assumes no more than two uniquely mapped segments and one breakpoint per split read^22^, *Picky* incorporates an algorithmic rationale to interpret long reads spanning multiple breakpoints that may even encompass the entire SV. Therefore, *Picky* has sufficient sensitivity to capture small SVs, particularly short tandem duplications. For example, a 0.9-Kb tandem duplication on chr.10p12.31 was misclassified as translocation by LUMPY but uncovered by *Picky* from a 12.9 Kb nanopore read (**Fig. 2c**). This duplicated region is a dinucleotide (TA)_n_ microsatellite repeat known to be highly variable in the human genome^23^. Despite its highly repetitive nature, this 0.9-Kb tandem duplication can be detected and accurately mapped due to the alignment specificity offered by the long read and the algorithmic design of *Picky*.

To confirm the accuracy of *Picky*, we validated SVs via PCR across either identified breakpoint junctions (TLCs, INVs and TDJs) or full-length rearranged regions (INSs, DELs and TDCs). All together, we validated over 200 SV events based on the presence of PCR-amplified products of the expected sizes indicated the positive validation status (**Supplementary Table 4**). 100% (40/40) of the predicted SVs that were supported by more than one nanopore read were validated, while 136 out of 173 (79%) SVs that were supported by only a single nanopore read were validated (**Fig. 3b** and **Supplementary Fig. 6a**). Using the 176 validated SVs as the reference data set, we further compared the specificity between nanopore long read and short read analyses. As shown, short-read sequencing analysis by LUMPY could only accurately detect a subset of SV classes (TLC, TDJ and DEL) given sufficient coverage (>30X), but drastically underperformed in classifying short-span TDs (TDC) and inversions, and was unable to unambiguously define insertion events (**Fig. 3c**). To further evaluate the sensitivity of *Picky*, we examined whether it could detect and classify the SVs identified by a previous short-read data analysis^20^. Excluding from the 27 SV regions not covered by nanopore reads, all the remaining 69 SVs were correctly classified by *Picky* to their corresponding types (**Supplementary Fig. 6b**). Selective examples of the validated SVs and their breakpoint junctions are shown in **Fig. 4**. This confirms that nanopore long reads, combined with the *Picky* analysis pipeline, enable accurate genome alignment, resolve the organization of repetitive sequence regions, and are clearly superior to short read approaches in identifying a wide range types of SVs.

**Figure 4.**
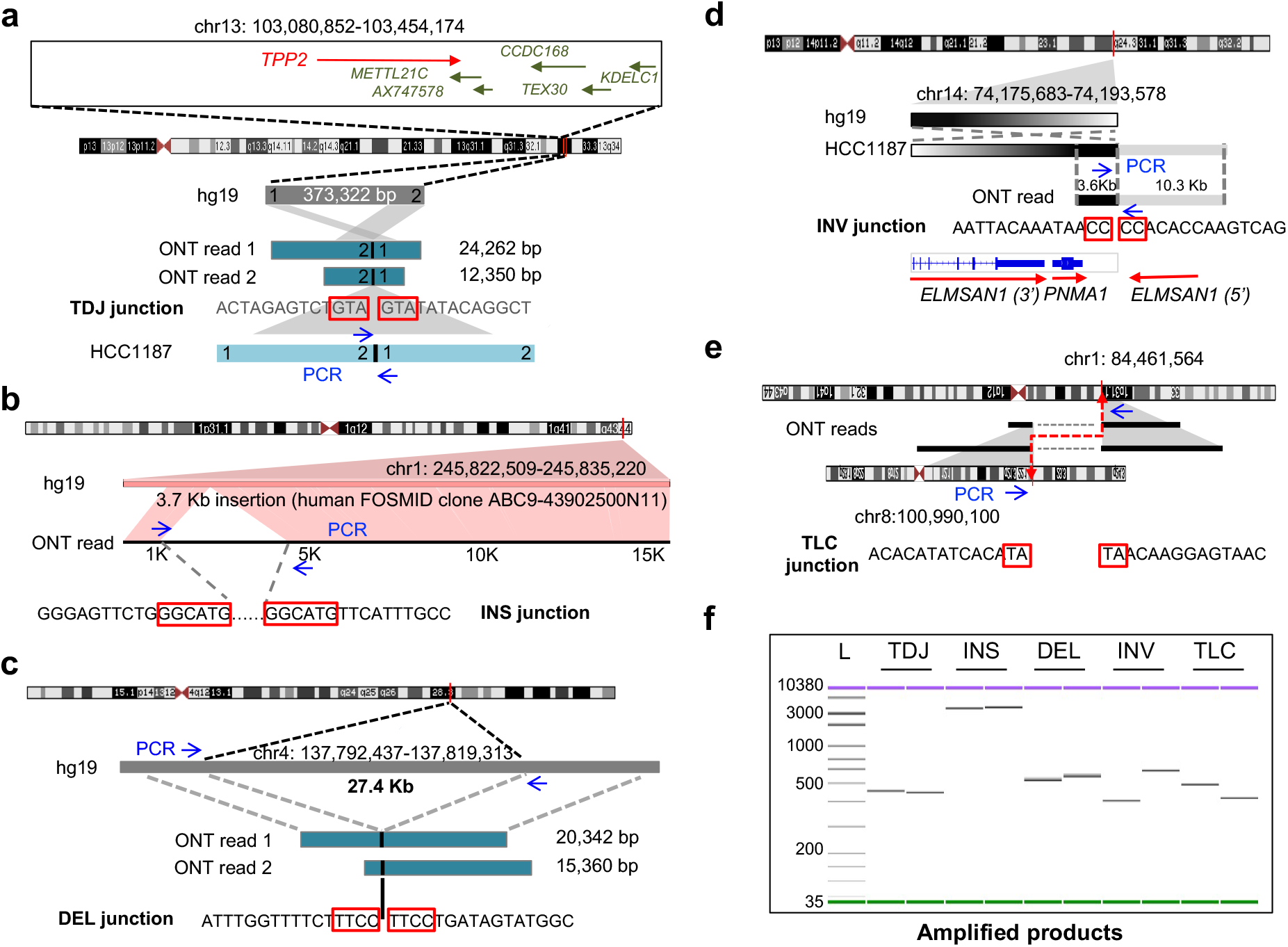
Examples of validated breakpoints and their detailed junction sequences. Nanopore read-to-genome alignments, junction sequences and affected genes were shown in each SV class. The micro-homologous sequences shared between junctions were highlighted in red boxes. (**a**) TDJ. (**b**) INS. (**c**) DEL. (**d**) INV. (**e**) TLC. The translocation t(1;8) identified is consistent with translocation identified previously by spectral karyotyping (SKY)^27^ with base resolution. (**f**) Amplified PCR fragments across breakpoints for each SVs shown in (**a**)-(**e**) were analyzed by Bioanalyzer (Agilent Technologies). L: molecular size markers.

### Nanopore long reads uncovered short-span SVs enriched in repetitive DNA sequences

SVs identified in HCC1187 genome exhibit a broad span distribution, ranging from 20 bp up to 100 Mb (**Fig. 5a** and **Supplementary Fig. 7**). Comparing to the short-read analysis, *Picky* uniquely detects short INSs, DELs, INDELs and TDCs through nanopore long reads spanning across the entire variable regions as evidently from their span size distribution. Vast majority of the 28,402 INSs and DELs and 3,332 TDCs span less than 1Kb with notable peaks around 300 bp (**Fig. 5a** and **Supplementary Fig. 7**), suggesting that they are enriched with repetitive sequences. When we examined their repeat content, simple INSs, DELs and TDCs exhibited a bimodal distribution based on the fraction of the SVs overlapping with repetitive sequence regions. 59-60% of the TDCs, INSs and DELs have more than 75% of their SV regions overlapping with repetitive sequences compared to 40% of the randomly selected genomic control regions of similar size distribution have more than 75% of their spans overlapping with repeats (**Fig. 5b**). Further dissecting the types of repeats found across different SV types, we found that the SINE and simple repeats are the predominant enriched repeat classes (**Fig. 5c** and **Supplementary Fig. 8a,b**). Therefore, small-size INSs, DELs and TDs account for the majority of the SVs detected in HCC1187 genome and they are predominantly copy number variation in repetitive sequence regions.

**Figure 5.**
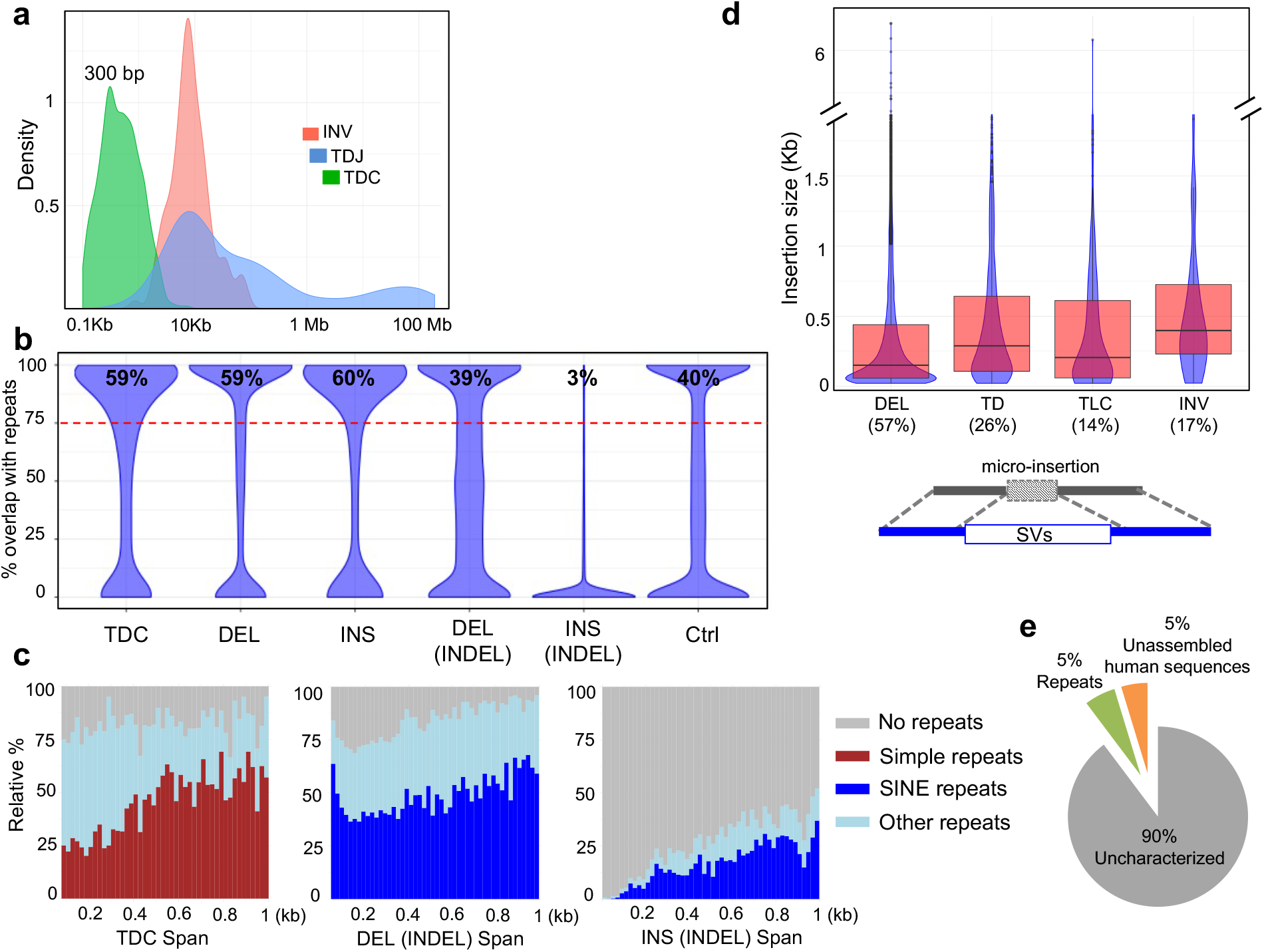
Long reads uncover repeat-rich SVs and the presence of micro-insertions within SV junctions. (**a**) Span distribution of INV, TDJ and TDC. **(b**) Fraction of regions overlapped with repeats among different SV classes. **(c**) Relative percentages of the major repeat types across different span sizes. **(d**) The percentages of SVs with micro-insertions and the size distribution of the inserted sequences. (**e**) The identity of the micro-insertions.

### Micro-insertion is a prevalent structural feature found within breakpoint junctions

Nanopore split reads aligned across SV junctions enable us to characterize breakpoint junctions in their entirety at nucleotide-resolution. When we closely examined the 66,660 breakpoint junctions from different SV types, we observed the presence of additional non-aligned DNA sequences inserted within the breakpoints. While these insertions were most frequently detected within the junctions of DELs (57%) and TDs (26%), they were also found in a subset of TLCs (14%) and INVs (17%) (**Fig. 5d**). The majority of these inserted sequences spans less than 500 bp, although some of them can be as long as 6 Kb, and has no significant similarity to the reference human genome. BLASTing these insertions against the NCBI non-redundant nucleotide database reveals that 90% of the inserted DNA pieces are completely novel with no homology to any known sequences, 5% of them overlap with known repetitive elements (mostly SINE/Alu), and 5% of them came from un-assembled human sequences, including newly emerged retrotransposed pseudogenes (**Fig. 5e** and **Supplementary Fig. 9a**). The small size of these sequences and their lack of significant homology with any existing known sequences, are consistent with the “genomic shards” observed in a few selective rearrangement events^24, 25^.

Formerly reported as rare events, these micro-insertions are thought to derive from non-templated DNA synthesis at the rearrangement junctions between chromosomal double-strand breaks^20^. Besides the micro-insertions, short stretches (usually 2-6 nucleotides) of identical sequence, known as overlapping microhomology^26^, were frequently spotted at SV breakpoint junctions (highlighted in **Fig. 4** and **Supplementary Fig. 9b**). Despite the visual appearance of such micro-homologies, nanopore reads currently do not have sufficient nucleotide-level accuracy to systematically characterize its exact context and frequency within the breakpoint junctions.

In summary, nanopore long-read based SV analysis uncovers large number of variants in repetitive sequence regions and precisely characterize breakpoint sequences. The micro-insertions of novel sequences associated with breakpoint junctions suggest that *de novo* DNA synthesis is a potential mechanism used in the non-homologous end-joining (NHEJ) during genomic rearrangement repair process.

### Breakpoint landscapes is associated with chromatin organization and transcriptional regulation

The distribution of the large numbers of SVs reflects the highly jumbled nature of HCC1187 genome (**Fig. 6a**). From the 1,911 TLCs, high frequency translocations between t(2;19), t(2;17), t(1;8) and t(10:13) were consistent with the translocations previously described by SKY (spectral karyotyping)^27^. The density of breakpoint distribution across the genome was found to associate with the fraction of genome annotated with gene coding regions. Using 1 Mb genomic span as the bin size, 80 out of 107 (75%) of the gene-rich regions (>10% gene-annotated nucleotides) have ≥ 20 breakpoints; while 180 out of the 193 (93%) gene desert regions (zero annotated nucleotide) have ≤ 6 breakpoints. Across genomic regions with different breakpoint densities, the top 10% breakpoint dense regions (> 40 breakpoints /Mb) have a significantly higher percent of nucleotides coding for genes than the bottom 10% breakpoint poor regions (< 9 breakpoints/Mb) (p < 2.2e-16, Mann-Whitney test) (**Fig. 6b**); suggesting high transcription activity could impact genome fragility. Interestingly, two of the hyper-density breakpoint loci (2q21.3–2q22.1; 65 breakpoints/Mb and 4q35; 139 breakpoints/Mb; highlighted in **Fig. 6a**) had very high inter-chromosomal contact probability (ICP) values (within top 10 percentile of ICP) found in the GM12878 cell line. ICP, defined as the sum of a region’s inter-chromosomal contact frequencies divided by its total contact frequencies from a nuclear ligation assay (Tethered Conformation Capture), has been used to describe the propensity of a region to form inter-chromosomal contacts within interphase nucleus^28^. The 4q35 subtelomeric region (chr4: 190,466,668-190,854,960) locates at nuclear periphery^29^ and 2q21.3–2q22.1 (chr2: 132,839,065-133,225,936) lies next to the vestigial centromere^30^. This raises the possibility that chromosome regions with frequent exposure to other chromosome and/or residing at the exterior of a chromosome territory, could be more prone to chromosomal breaks compared to the protected regions deeper inside the chromosome territory. Further analysis on the whole genome level indicated a positive correlation between ICP and breakpoint count (**Fig. 6c**). For instance, the group with the highest breakpoint counts (> 30) exhibits higher ICP values (p-value < 1.6e-16, Mann-Whitney one-sided non-parametric test) than the group with the lowest breakpoint counts (< 10). These observations further support the notion that intermingling of chromatin organization directly influences the structural properties associated with elevated frequency of DNA double strand breaks^31, 32^.

**Figure 6.**
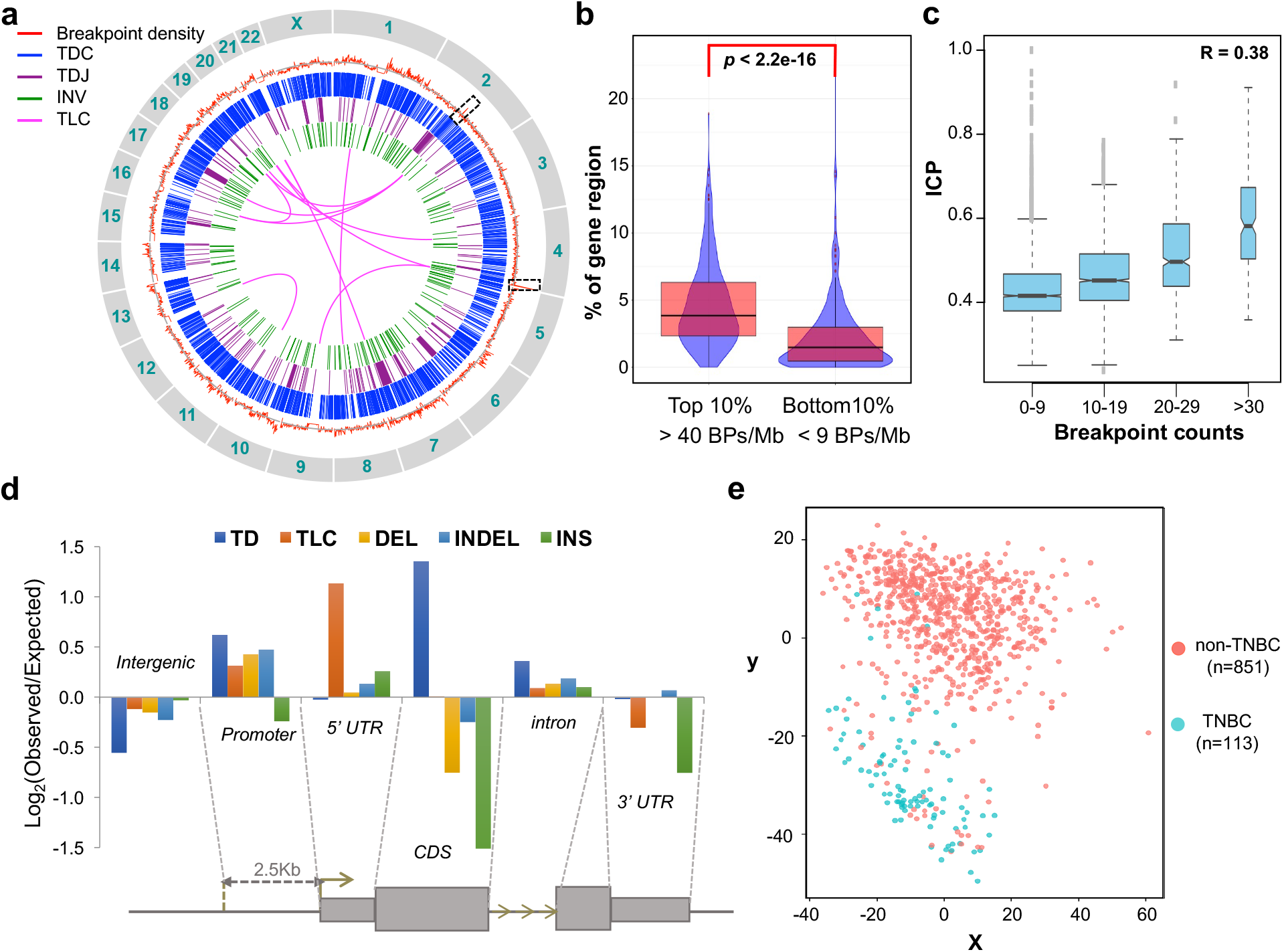
The analysis of genomic distribution of breakpoints and their affected genes. (**a**) Genome-wide distribution of the SVs and their associated breakpoints density. From outer circle to inner circle: Red: measured breakpoint density; Grey: reference average density of 22 breakpoints/Mb; Blue: TDC; Purple: TDJ (with span size < 20 Mb); Green: INV; Magenta: TLC (with pair counts ≥ 4) at 1Mb bin size. **(b**) Fraction of regions with transcribed genes in regions of the top and bottom 10% breakpoint densities. BP, breakpoint. *P* value was calculated by Mann-Whitney test. **(c**) The trend of increasing ICP with increasing number of breakpoints. In the plots, the breakpoint and ICP were calculated for every chromosome segments of 138 HindIII sites (**Methods**). The ICP values were then grouped based on the breakpoint counts as indicated on the x-axis. The R value is the Pearson’s correlation coefficient on the underlying, ungrouped data. (**d**) Enrichment of breakpoint from each SV class distributed along different genomic features. (**e**) The multidimensional scaling (MDS) plot of SV-genes in the BRCA dataset within the TCGA.

### Enrichment of SVs in the regulatory repertoire of the genome with impacts on gene expression

The high frequency of SVs associated with the interconnected chromatins of high gene density further implicates linkage between genome structure and transcriptional regulation. Thus, we investigated the distribution of different SV associated-breakpoints among intergenic, coding sequence (CDS), promoter (2.5 Kb upstream of TSS), 5’ and 3’ un-translated regions (UTRs), and introns (**Fig. 6d**). Significant enrichment of breakpoints was found in promoters and 5’ UTR for most of the SV types, particularly the TLC and TDs. In contrary, CDS and 3’ UTR were largely depleted of most of the SVs except TDs. Such patterns suggest that the potential impact of SVs is to modulate gene transcription by altering regulatory elements in promoters and 5’ UTR while minimizing their detrimental effects on functional protein integrity or RNA stability. Interestingly, repeat-rich versus repeat-poor TDs exhibited contrasting distribution patterns (**Supplementary Fig. 10**). Breakpoints in the repeat-poor TDs generated by NHEJ or microhomology mediated end joining (MMEJ) were found at higher frequency in all compartments of gene regions, particularly in CDS. This is consistent with previous findings of short TDs (Tandem Duplicator Phenotype; TDP) in TNBC, ovarian cancer, and endometrial carcinoma where tumor suppressor genes statistically more frequently disrupted than random^10^. This strongly suggests that a specific SV, tandem duplications, generated by different repair mechanisms, homologous recombination (HR) or NHEJ/MMEJ, have different functional consequences in cancer biology.

SVs occurring in promoters/regulatory elements can selectively lead to oncogene activation or tumor suppressor gene inactivation. Such effects on gene expression are likely to be cancer pathway or origin specific. To test this, we determined the genes affected by the SVs and examined their expressions from 113 TNBC and 851 non-TNBC tissues of the breast carcinoma (BRCA) dataset within the cancer genome atlas (TCGA)^33^. In contrast to the control genes and with permutation, the expression of 1,260 coding genes disrupted by the major SV classes (INDEL and TD) effectively distinguished TNBC from non-TNBC types as visualized by multidimensional scaling (MDS) plot (**Fig. 6e** and **Supplementary Fig. 11**), highlighting the functional impacts of SV analysis and its link to tumor molecular classification.

## Discussion

Existing short read sequencing and SV analysis tools are limited in resolving complex structural variation and delineating molecular structures of breakpoints, particularly within repetitive regions. Here, we leverage the recent technology advancement of nanopore sequencing and a customized long-read specific SV analysis pipeline to detect a full spectrum of structural variants in a highly complex cancer genome model. Our results demonstrate that nanopore sequencing offers superior performance to high depth, short-read analysis in specificity for SV detection, even at a modest depth. The specificity of the long read-to-genome alignment uncovers short span SVs residing in different repetitive elements, which are highly abundant but largely understudied by existing approaches. Repetitive elements are known to provide architecture framework in genome organization^34^ and as a major source of variation inducing human diseases^35^. Missing SVs localized to these repetitive regions severely limits the analysis of important structural mutations in cancer, especially those SVs in cancers with intact homologous recombination mechanisms.

Our breakpoint analysis suggests the linkage between the propensity of inter-chromosomal connectivity and frequency of genomic lesions. Nuclear regions with high transcriptional activity are shown to have extensive inter-chromatin contacts^28^ and intermingling between chromatin territories can influence translocation efficiency^31^. Therefore, the accessibility and conformation in the active chromatin domains may provide the structural basis of genome fragility. The preferential enrichment of breakpoints in the regulatory repertoire of the genome further implicates a potential mechanism in which genome rearrangement can reconfigure the transcriptional program in the cancer transformation process. Expanding the high-resolution of SV analysis throughout tumorigenesis should further our understanding of the homeostasis between genome architecture, variation and carcinogenesis.

Long-read sequencing possesses many unique features that improve from the current state of SV detection. Yield from nanopore has been dramatically increased for the past year with higher sequencing speed and pore stability. Furthermore, multiple strategies have been devised in parallel to improve the basecalling accuracy, including a new basecaller Scrappie^36^. With the continuous progress made on throughout, accuracy and analytic tools, genome-wide haplotype specific SV detection can be attempted to interrogate the diversity and complexity of cancer genome variation. Given the superior alignment specificity, higher resolution and broader utility of single molecule long-read data, we anticipate that there will be soon a paradigm shift on the sequencing approaches for genome structural analysis; which will reveal new insights in the dynamics of SVs and the mechanisms of their generation during tumorigenesis.

## Methods

### Nanopore long read sequencing

High molecular weight DNA was extracted from HCC1187 cells by MagAttract HMW DNA Kit (Qiagen, 67563) followed the manufacturer’s instructions. Briefly, 1 × 10^6^ frozen cells were lysed with 220 μL of Buffer ATL, 20 μL Proteinase K and incubated overnight at 56°C with 900 rpm. 4 μL RNase A was added to cleave RNA. 150 μL Buffer AL, 280 μL Buffer MB and 40 μL MagAttract Suspension G beads were then added to capture the HMW DNA. Next, the beads were cleaned up by 700 μL Buffer MW1, Buffer PE and NFW followed by elution with 150 μL Buffer AE. Nanopore sequencing libraries were prepared according to the target size and the sequencing kits supplied by Oxford Nanopore Technologies (ONT) (**Supplementary Table 5**). HMW genomic DNA was fragmented by either miniTUBE Blue (Covaris, 520065, for 3 Kb), miniTUBE Red (Covaris, 520066, for 5 Kb) or g-TUBE (Covaris, 520079, for 8 Kb and 12 Kb). For libraries targeted at 12 Kb, size selection was performed for sheared fragments larger than 10 Kb using 0.75% agarose cassette (Sage Science, BLF7510) by Blue Pippin^TM^ DNA Size Selection System. For libraries of less than 10 Kb, AMPure XP beads (Beckman Coulter, A63881) were used for clean-up. Next, libraries were prepared according to recommendation by ONT. Briefly, NEBNext FFPE RepairMix (NEB, M6630) was added to repair nicks in the DNA. Then end-repair and dA-tailing were performed using NEBNext Ultra II End-Repair/dA-tailing Module (NEB, E7546). For 2D libraries (WTD01-WTD13), we prepared the ligation reaction as below: 38 μL water (DNA), 10 μL Adapter Mix (AMX), 2 μL Hairpin Adapter (HPA) and 50 μL Blunt/TA Ligase Master Mix (New England Biolabs, M0367). Ligation was performed at room temperature for 15 minutes. 1 μL Hairpin Tether (HPT) was added to the reaction and incubated at room temperature for 15 minutes. Then 50 μL MyOne C1 beads (Thermo Fisher, 65001) beads were prewashed twice with 100 μL Bead Binding Buffer (BBB). The MyOne C1 beads resuspended in 100 μL BBB were added to the ligated DNA reaction and incubated on a rotator at room temperature for 15 minutes. The beads were washed twice with 150 μL BBB and eluted in 25 μL Elution Buffer (ELB). For 1D libraries (WTD14-WTD15), we prepared the ligation reaction as below: 30 μL water (DNA), 20 μL Adapter Mix (AMX) and 50 μL Blunt/TA Ligase Master Mix (New England Biolabs, M0367). Ligation was also performed at room temperature for 15 minutes. The AMPure XP beads were resuspended at room temperature by vortex. 40 μL of beads were added into the ligation product. The beads were washed twice with 140 μL Adapter Bead Binding (ABB) buffer and eluted in 25 μL Elution Buffer (ELB). The eluted product was the adaptor-ligated library as the Pre-sequencing Mix (PSM) used in nanopore sequencing.

The libraries were sequenced on MinION Mk1b device (ONT) using R9 and R9.4 flow cells following the standard 48 h run scripts (**Supplementary Table 5**). Real-time basecalling was performed on EPI2ME cloud platform (ONT). Read sequences were extracted from base-called fastf5 files by Poretools (version 0.5.1) to generate fastq file. All 2D reads from WTD01-13 (2D ligation libraries) and all 1D reads from WTD14 and WTD15 (1D ligation libraries) were used for subsequent analysis.

### PacBio sequencing

DNA templates were prepared as described in 12 Kb nanopore sequencing libraries. A PacBio genomic DNA library was prepared (12 Kb) according to the manufacturer’s instruction (Greater Than 10 Kb Template Preparation Protocol). The library was sequenced on PacBio RS II instrument. PacBio sequencing data was processed by the PacBio SMRT Portal pipeline of Read of Insert with the parameters “min full pass = 0” and “min predicted accuracy = 75”.

### Picky pipeline for SV detection

The reads (in fastq format) were processed with an in-house assembled analysis pipeline *Picky* (**Supplementary Fig. 12**). Briefly, *Picky* is composed of three steps: aligning nanopore reads to the reference genome, picking the best alignments and calling structural variants (SVs). Firstly, the nanopore reads were mapped to human genome (hg19) using the LAST aligner (last755)^21^ with the parameters “-r1 -q1 -a0 -b2”. For high sensitivity, we adopted the scoring scheme (reward = 1, penalty = −1, gap open = 0, gap extension = 2) used in NCBI megaBLAST (https://www.ncbi.nlm.nih.gov/books/NBK279678/). LAST aligner reported all high scoring pair (HSP) segments for each nanopore read. Next, segments with EG2 > 1.74743e-12 or %Identity < 55 were discarded. Nanopore reads with no aligned segment post-filtering were considered unmapped and reads with just a single segment were reported as uniquely mapped. For read with multiple segments, the best segment (lowest EG2 and highest score) was used to determine the EG2 threshold (EG2_best_) and %identity (%identity_best_). Segments which EG2 < 1e-49 and the absolute difference between alignment %identity and %identity_best_ <= 10% were retained. To speed up processing, segments were grouped into representative segment groups based on the read coordinates. Specifically, a representative segment group may contain three subgroups, namely ‘exact’, ‘subset’, ‘similar’. Segments in subgroup ‘exact’ share the exact read start and end coordinates although their genomic coordinates differ. Subgroup ‘subset’ contains segment where its read coordinates are just a sub-span of the representative segment for the segment group. Members of subgroup ‘similar’ have 95% reciprocal overlap and %identity difference <= 5 with respect to the group’s representative segment. A greedy seed-and-extend strategy was used to stitch segments together and produce the combined alignment that spans the most of the nanopore read. The segment groups which EG2 = EG2_best_ were designated as the seeds. If there were >= 5 seeds, we used the top 200 of them as potential extenders. Otherwise, potential extenders could consist of the segment group(s) where EG2 = 0 minimally, with the successive inclusion of the 4 segment group collections binned by 0 < EG2 <= 1e-300, 1e-300 < EG2 <= 1e-200, 1e-200 < EG2 <= 1e-100, and 1e-100 < EG2 <= 1e-50, as long as the total number of potential extenders did not exceed 100. The algorithm started by picking a seed and attempted to extend it with the potential extenders to maximize the coverage of a nanopore read. For an extender to be chosen it must extend the current read coverage by at least 200 bases and the overlap between the extender and current combined segments must account for < 50% of each length. This process then continued with new combined segments until the whole nanopore read was covered or we run out of candidate extender. We then scored all combined alignments (the permutation of 5’ extensions, seed and 3’ extensions) as the sum of segments’ score and discarded any “combined alignments” that did not cover more than 70% of the read. All combined alignments with score within 90% of the best score for a nanopore read were reported.

If no combined alignment can be derived, this nanopore read was classified as “without candidate”. If multiple combined alignments remained, it was classified as multi-candidates read. When only a single combined alignment remains, it may consist of a single HSP (SCSF) or multiple HSPs (SCMF). SCMF reads were used to capture potentially SV(s). SV calling was performed on SCMF where each HSP mapped to a single locus (SCMFSL, aka split-read) while SCMF with HSP(s) mapping to multi-loci (SCMFML) was ignored. Finally, SV calling was performed with an in-house script based on the order and the distance between split-read’s aligned segments. This distance was defined as sDiff when computed with the reference genome coordinates and as qDiff when the reads coordinates was used. Based on this information, SV was called as follow: Insertion (INS): split-reads with sDiff < 20 and qDiff > 20. Deletion (DEL): split-reads with sDiff > 20 and qDiff < 20. Insertion co-occurred with deletion (INDEL): split-reads with sDiff >= 20 and qDiff >= 20. Inversion (INV): split-reads from same chromosome and with different orientation. Translocation (TLC): split-reads from different chromosomes. Tandem duplication detected by junction split-reads (TDJ): split-reads only captured the duplication junction (tail and head of the duplicated fragment). Tandem duplication detected by complete duplicated segment (TDC): split-reads captured the complete duplicated fragment and sDiff <= −100. Full list of SVs from each of the seven SV types can be found in **Supplementary Table 3**. *Picky* is also conscious of the nanopore high error rate in homopolymer sequences and implements a correction mechanism to reduce its impact on SV detection. Specifically, we observed that homopolymers in all four nucleotide contexts (A_n_, T_n_, C_n_ and G_n_) were significantly underrepresented relative to expectation (**Supplementary Fig. 13a**). Examining the raw signal data and their corresponding called bases, we concluded that homopolymers beyond five identical bases are not correctly reported and lead to false-positive classification as deletions (**Supplementary Fig. 13b**). Therefore, we further adjusted *Picky* to ignore deletions found in the homopolymer regions. As a result of this adjustment, the numbers of SVs classified as insertions, deletions or INDEL decreased from 29,977 to 28,402.

### Comparison between long read and short read data

Nanopore reads aligned uniquely (possibly multiple fragments) to the human genome (hg19) were extracted for genome coverage computation with BedTools (v2.25.0). Similarly, HCC1187 Illumina paired-end sequencing data (SRA accession SRX969058) was mapped to human genome with BWA-MEM (v0.7.12). The mapped reads were sampled at 2.5X, 10X, 30X and 60X. Genome coverage was computed on reads with a mapping quality of 60.

### Phased adjacent SVs from multi-breakpoint long reads

We counted the pair of adjacent SVs called in all the multi-breakpoint long reads. We assumed that the expected count would follow the distribution of independently drawing two SVs from the population of the SVs from all the multi-breakpoint long reads. The log-likelihood was then computed.

### SVs span size distribution

To explore the genomic feature of SVs, we applied different methods to determine the distributions of their span size. For DEL, INS and INDEL, we calculated the total numbers of SVs from each genomic bin (bin size = 20 bp). For INDEL, we used sDiff and qDiff as deletion and insertion spans, respectively. For TDC, TDJ and INV, density plots were generated from their span distributions to show their large size variations.

### Characterization of genomic features associated with breakpoints

Genomic distribution of the SV and the breakpoint Breakpoint density was computed from the numbers of breakpoints per Mb across the genome. The density of TLC pair was calculated using the TLC breakpoint distribution in pair per Mb across the genome. Circos plot was performed using the breakpoint densities, spans of TDC, TDJ (< 20 Mb), INV and TLC pairs (counts >3).

### Association between gene coding and breakpoint density

Gene density was computed as the fraction of bases overlapping with annotated gene regions (exons and introns; GenCode V24) in each Mb bin across the genome. Violin plots of the gene density from the top 10% breakpoint dense regions (breakpoint density > 40) and the bottom 10% least dense regions (breakpoint density < 9) were generated. Mann-Whitney test was performed.

### Breakpoint landscape analysis

Each breakpoint was stepwise assigned to different class of genomic features based on the GENCODE v24 gene model (**Supplementary Fig. 14**). The promoter was defined as the upstream 2.5 Kb region of the transcription start site. The fraction of reference genome in each class was taken as the background distribution to compute the expected number of breakpoints for each SV type. For SVs with two breakpoints, the pair was considered independent. The ratio of number of breakpoints between observed and expected was log_2_ transformed.

### Repeat analysis

To determine whether the inserted DNA fragments from INS and INDEL as well as TD regions contained repetitive sequences, if so, which class of repeats, we extracted the inserted or duplicated DNA sequences from their corresponding nanopore reads and annotated them to different repeat classes by aligning them to public annotated repeat sequences using RepeatMasker-open-4-0-6 (http://www.repeatmasker.org/). Violin plot was generated with the percentages of the SV fragments annotated to repeats (hg19). The relative ratios of the most predominant repeat class, all other repeats and no repeat from each of the five SV types (TDC, deletion regions in INDEL, insertion regions in INDEL, DEL and INS) were produced in 20 bp span size intervals.

### Distribution of genomic micro-insertions

Un-aligned DNA sequences found between each breakpoint junction (nanopore reads alignment with qDiff > 20) were extracted from their corresponded nanopore reads from INDEL, TD, TLC and INV. Their size distribution was plotted for these 4 classes.

### ICP analysis

ICP was defined as the sum of a region’s inter-chromosomal contact frequencies divided by its total contact frequencies. We downloaded the TCC (Tethered Conformation Capture) interaction matrix from NCBI SRA accession SRX030110^32^ and computed the ICP on the whole genome^28^. The counts of breakpoints were partitioned into four groups from low to high. The correlation was plotted from ICPs found in different partitions.

### Multidimensional scaling of gene expression analysis

SVs with spans range from 1Kb to 1 Mb were selected to compare their impacts on gene expressions between TNBC and non-TNBC samples. There are 1,260 coding and 711 non-coding genes in the selected 537 DEL, 2,383 INDEL and 188 TDJ events (SV-genes). Their expression in 113 TNBC and 851 non-TNBC samples was retrieved from gene expression data of the breast carcinoma (BRCA) of the cancer genome atlas (TCGA) based on our previous study^33^. Two data sets, the control set and the permutation set, were used to evaluate the significance of grouping analysis. The control data was selected from the non-SV genes of equivalent number that were expressed at similar level as the SV-genes. For the permutation data, the expressions of SV-genes were sample-wise permutated to ensure that each gene has the same expression profile with a distinct sample order. Multi-dimensional Scaling (MDS) was used to analyze the expression of SV-genes in the BRCA dataset and visualize the sample relationship.

### Validation for SV candidates

We selected 3-46 SV events from each SV classification (**Supplementary Fig. 6**). For candidates in DEL, INDEL, INV, TLC and TDJ, we designed primers to detect the breakpoints. For candidates in INS, we designed primers to amplify the entire inserted fragments. For candidates in TDC, we attempted the validation of the breakpoints on 13 candidates and the validation of the entire duplicated regions on 35 candidates. All primers used in this study are provided in **Supplementary Table 6**.

### Checking of short reads called SV against PCR validated SV

HCC1187 Illumina paired-end sequencing data (SRA accession SRX969058) was mapped to human genome (hg19) with BWA-MEM (v0.7.12). The mapped reads were also sampled at 2.5X, 10X, 30X and 60X. LUMPY (v0.2.13) was used to call SVs using both non-redundant split-read and discordant paired-end reads with the minimum weight for a call (-mw) as 2, 3, 5, 10, and 16 for the subsampled sets 2.5X, 10X, 30X, and 60X and the whole dataset (102X) respectively. We then loaded the LUMPY generated vcf files in IGV browser to visual check the PCR validated SVs locus for the same SV type called by LUMPY.

## Acknowledgments

The authors thank P. Shreckengast for collecting the HCC1187 cells; and C. Robinett and A. Lau for their comments on the manuscript; and B. Hanson and M. Bolisetty for their help in setting up our initial nanopore runs. Research reported in this publication was partially supported by the National Cancer Institute of the National Institutes of Health under Award Number P30CA034196.

## Competing financial interests

L.G., C.-H.W. and C.-L.W. have received a few batches of reagent from Oxford Nanopore. C.-L.W. has received travel and accommodation support from Oxford Nanopore as an invited speaker at the Oxford Nanopore user meeting.

## Supplementary Information

**Supplementary Figure 1. Performance of the nanopore long read data.** (**a**) Yield of 15 nanopore runs of different chemistry versions. (**b**) Accuracy and read counts from runs with different run protocols and pore speeds. Grey, 2D and R9 (250 bases/sec). Red, 2D and R9.4 (250 bases/sec). Blue, 1D and R9.4 (450 bases/sec). (**c**) Genome coverage from the 7.9 Gb of the nanopore data generated in this study.

**Supplementary Figure 2. Comparison of the genome coverage from long read and short read data.** (**a**) The percentage of the genome covered by short versus long read data of different depth. (**b**) A region (chr1: 25,732,083-25,735,365) covered by a 14.7-Kb nanopore read; which remains a gap even from 185 Gb, 60X of short-read data. This region is rich in SINEs/LINEs.

**Supplementary Figure 3. Analysis of read length bias from nanopore and PacBio reads.** (**a**) Length distribution of nanopore reads from 12 Kb target template size (data from WTD03, WTD07–WTD12). (**b**) Length distribution of nanopore reads from 3 Kb target template size (data from WTD04–WTD06). (**c**) Length distribution of PacBio reads from the same 12 Kb target template size.

**Supplementary Figure 4. The relationship between the percentage of split-reads and the mean read length.**

**Supplementary Figure 5. Analysis for phased SVs by the multi-breakpoint long reads.** (**a**) The total counts and (**b**) the log-likelihood of the adjacent SVs phased by the multi-breakpoint long reads. Red count indicates observation > 2x expected. Blue count indicates observation < 0.5x expected.

**Supplementary Figure 6. The sensitivity and specificity of the *Picky*-called SVs.** (**a**) Summary of the validated SVs by PCR strategy. (**b**) The numbers of high confidence SVs previously described in HCC1187 detected by nanopore sequencing.

**Supplementary Figure 7. The span distribution of DEL, INS and INDEL.**

**Supplementary Figure 8. SVs enriched in repeat regions.** (**a**) Relative percentages of repeats across different span sizes in simple DEL. (**b**) Relative percentages of repeats across different span sizes in simple INS.

**Supplementary Figure 9. Selected cases of validated SVs.** (**a**) An insertion of 1.7 Kb *CBX3* retrotransposed pseudogene in chromosome 15 revealed by a 24.8 Kb read (WTD13:1125). (**b**) A confirmed TDJ of 1.87 Mb on chromosome 18 revealed by a 3.0 Kb read (WTD02:4697).

**Supplementary Figure 10. Distribution of the breakpoints associated with TDC along different genomic features of transcription.**

**Supplementary Figure 11. Control of the multidimensional scaling (MDS) analysis.** (**a**) Histogram of gene expression from SVs-genes (log_2_ transferred). (**b**) Histogram of gene expression from the control genes. Similar expression profiles and the equivalent numbers of SVs-genes are shown (log_2_ transferred). (**c**) The MDS plot expressions of the SVs-genes by sample-wise permutation. (**d**) The MDS plot of the expressions from the control genes. All data are from the breast carcinoma (BRCA) dataset within the cancer genome atlas (TCGA).

**Supplementary Figure 12. Overview of *Picky’*s process and categories assigned for a nanopore read.**

**Supplementary Figure 13. Homopolymer analysis of nanopore reads.** (**a**) The ratio of the observed versus expected instances of all 1,024 5-mers. Highlighted are the 4 under-called homopolymers. (**b**) The annotated current trace for the segment harboring basecalled deletion. The trace indicates the clear existence of the two homopolymers (marked (A)_20_ and (T)_18_) rather than the deletion flanked by (A)_5_ and (T)_5_.

**Supplementary Figure 14.** Overview of the process assigning breakpoints to their corresponding genomic features based on the gene model.

**Supplementary Table 1.** Summary of the 15 nanopore runs in this study.

**Supplementary Table 2.** Summary of the mapping and the SV calling results.

**Supplementary Table 3.** List of seven SV types detected in nanopore data.

**Supplementary Table 4.** SVs selected for validation analysis.

**Supplementary Table 5.** Details of nanopore sequencing kits, devices and software.

**Supplementary Table 6.** List of all primers used in this study.

